# WHOOPER Web application for Hands-On identification of proteins co-Occurrence among Phyla, focused on user ERgonomics

**DOI:** 10.1101/2025.06.19.660341

**Authors:** Sylvain Marthey, Natacha Baffo, Véronique Martin, Laiqa Zia Lodhi, María-Natalia Lisa, Gwenaëlle André

**Affiliations:** Université Paris-Saclay, INRAE, MaIAGE, Jouy-en-Josas, France; Instituto de Biología Molecular y Celular de Rosario (IBR CONICET-UNR), Ocampo y Esmeralda, Rosario S2002LRK, Argentina; Université Paris-Saclay, INRAE, BioinfOmics, MIGALE bioinformatics facility, 78350, Jouy-en-Josas, France; Plataforma de Biología Estructural y Metabolómica (PLABEM), Ocampo y Esmeralda, Rosario S2002LRK, Argentina

## Abstract

**Summary:** WHOOPER is an intuitive web application designed to facilitate the analysis of protein co-occurrence across taxa. By integrating HHblits and HMMER, it enables efficient detection of remote homologs across extensive proteomic datasets. Its results interface combines an original data representation with advanced dynamic filtering functionalities to enable real-time exploration and exploitation of alignment data. Addressing key limitations of existing tools, WHOOPER streamlines workflows for studying protein prevalence, co-occurrence, and phylogenetic patterns, supporting seamless export for downstream evolutionary and functional genomics research.

**Availability and implementation:** The code for Whopper is available at https://github.com/SmartBioInf/whooper. A public instance allowing the screening of more than 7500 reference proteomes from eukaryotes, bacteria, and archaea is available at: https://whooper.migale.inrae.fr

## 1. Introduction

Understanding the evolution of biological functions across different taxa requires evaluating the distribution and prevalence of the proteins involved in specific pathways, as well as examining their co-occurrence and co-evolution within genomes. The increasing availability of genomic data makes it now possible to study gene co-occurrences on a broad taxonomic scale, thereby revealing evolutionary events and their correlation with phenotypic traits (Pende *et al*, 2021 [1]; Martinez *et al*, 2023 [2]). This type of research not only requires the accurate identification of homologous proteins across diverse phylogenetic distances, but also requires comparing the prevalence of each protein relative to others, in conjunction with their position on the phylogenetic tree.

Several online applications allow for the identification of protein homologs across large reference proteomes (see Supplementary Table 1). This identification can be (i) direct, through homology searches based on sequence (NCBI Blast, Boratyn *et al*, 2013 [3]; UniProt, UniProt Consortium, 2023 [4]; webFlaGs, Saha *et al*, 2021 [5]; Annoview, Wei *et al,* 2024 [6]), profile (fast.genomics, Price *et al*, 2024 [7]; MPI Bioinformatics Toolkit, Gabler *et al*, 2020 [8]; HMMER web server, Potter *et al*, 2018 [9]), or three-dimensional structure (Foldseek, van Kempen *et al* 2024 [10]) across all proteins in a dataset. Alternatively, identification can be (ii) indirect, based on the association between query sequences and their closest homologs retrieved from pre-computed datasets of potential orthologs and paralogs, using various methods and at different biological scales; this association can be explored through protein function (InterProScan, Jones *et al,* 2014 [11]; GeCoViz, Botas et al, 2022 [12]), protein-protein association networks (STRING, Szklarczyk *et al*, 2023 [13]), or phylogeny (MetaPhOrs 2.0, Chorostecki e*t al,* 2020 [14]; PhylomeDB 5, Fuentes *et al*, 2022 [15]).

Among these tools, some offer “Visual Analytics” capabilities (Pak Chung Wong, 2004 [16]), providing interpretable and interactive graphical representations of large datasets. This is achieved by grouping and contextualizing identified homologs alongside other relevant information, such as domain organization (HMMER, InterProScan), gene neighborhood (STRING, fast.genomics, webFlaGs, AnnoView, GeCoViz), or organism relationships, including phylogenetic data (NCBI Blast, UniProt, HMMER, fast.genomics, STRING, PhylomeDB 5, Foldseek) and habitat (GeCoViz). Despite these features, functionalities like dynamic filtering, selection, and exporting results of interest remain limited in most cases (Supplementary Table 1). This limitation is particularly problematic because individual results for each protein must be manually cross-referenced outside of these applications to conduct co-occurrence analysis. In fact, only two of them (STRING and fast.genomics) allow for the simultaneous analysis of multiple proteins, and only fast.genomics enables the identification of taxa without homologs. However, this one limits co-occurrence studies to just two proteins, while STRING does not provide a direct method for retrieving homologous sequences for specific taxa.

Despite the availability of web servers for remote homologs identification and proteins co-occurrence studies, there remains a need for a web application that: (i) is intuitive, interactive and user-friendly, (ii) enables fast and efficient identification of remote homologs on hundreds of proteomes, (iii) supports visual analytics of proteins prevalence and co-occurrence among phyla, (iv) facilitates the easy selection and export of the relevant results, and (v) is simple to configure for any set of proteomes. To address this gap, we developed WHOOPER, a web application that combines the remote homologs detection capabilities of HHblits and HHMMER with an interactive, ergonomic and innovative results interface. WHOOPER empowers users to quickly and efficiently analyze protein co-occurrence among phyla.

## 2. Features and methods / Software description

### 2.1 Research set up

On the submission interface, the search configuration is done by importing query sequences for alignment and selecting the relevant proteomes for screening. FASTA-formatted sequences can be added either by copy-paste or by uploading a file. Since proteome selection plays a crucial role in the analysis, several features have been implemented to help users quickly choose hundreds of them by species or entire taxonomic groups (Supplementary Figure 1): an autocomplete list, an interactive taxonomic tree, and the ability to import/export accession or taxon ID lists. The latter makes it easy to seamlessly iterate a new search based on the results of a previous analysis.

### 2.2 Homology searching

In the first step, for each query protein sequence, a Hidden Markov Model (HMM) profile is generated by aligning it against the UniRef clustered sequence database (Suzek *et al*, 2007[17]) using HHblits (Remmert *et al*, 2011 [18]). In a second step, these HMM profiles are aligned against all proteins from the selected species using the HMMsearch program from HMMER3 (Eddy SR, 2011 [19]). Combining these two tools enables the detection of distant homologs in the “twilight zone” of very low sequence identity, while preserving a straightforward and efficient storage system for reference genomes in FASTA format.

### 2.3 Results display, exploration and export

The result analysis environment is the core of Whopper, designed to streamline the rapid handling of results immediately upon generation. Following B. Shneiderman’s visual information seeking mantra (1996) “Overview first, zoom and filter, then details on demand”, this interface prioritizes ergonomics and user experience. The organization is set to ensure an intuitive and efficient workflow for exploring and interpreting protein co-occurrence and homolog data. It is structured into four key components (Figure 1): (1) Control Panel, allows users to configure and refine result filters interactively; (2) Hits Chart, provides an overview of the results, enabling users to visualize data trends and distributions; (3) Selection Table, displays selected results in a tabular format for easy review; and (4) Alignment Detail Display Area, offers detailed views of individual alignments for in-depth analysis.

**Figure 1.**
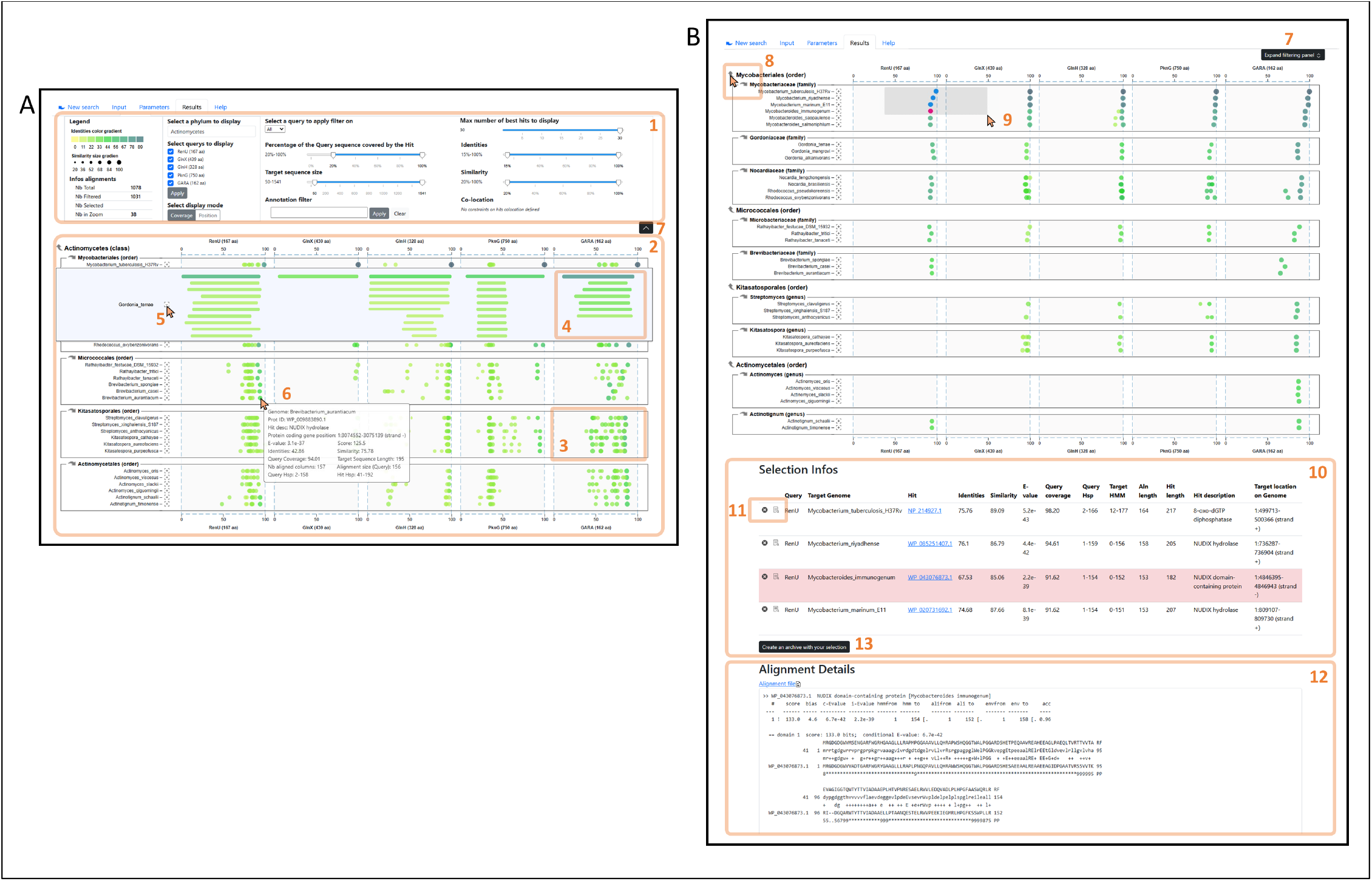
The WHOPPER browser with its key features. Here it is exemplified with PknG that cooccurs with proteins of its operon (GlnX, GlnH) and with its partner proteins GarA and RenU, with which it interacts in the regulation of glutamate metabolism (Bhattacharyya et al, 2018 [21]) and biofilm development (Wolff et al, 2015 [22]), respectively. (A) Raw alignments. (B) Filtered alignments. The control panel (1) includes display options and real-time filtering features, and can be collapsed or expanded (7) to maximize the results overview. The hits chart (2) enables visualization of all alignments for query sequences (on the X-axis) against all proteomes (Y-axis), grouped by taxon. Hits can be displayed in two ways: (i) points representing their coverage (3) or (ii) bars showing their position (4). The zoom feature (5) allows to display all the hits of a specific proteome. Hovering a hit displays its metadata (6). Taxonomic navigation (8) allows to change the taxon and its sub-taxons displayed. Hits are added to the selection directly on the graph by clicking on the hit to be added or by using a selection rectangle (9). The combination (*Ctrl* + *select*) enables to add/remove new selection to current selection. Selected hits are shown in a table (10), located below the hits chart, associated with their metadata. From this table, hits can be highlighted or removed from the selection. The alignment details of the highlighted hit are displayed below the selection table (12). A hit can also be highlighted by double-clicking it on the chart. The selection is available for download in different formats (13): fasta sequences grouped by query or genome, metadata with taxonomic information. The search parameters and the applied filters can also be downloaded.

The Control Panel, located at the top of the page, allows users to refine displayed results by limiting them to specific queries or taxa and dynamically filtering by various criteria. These criteria include: (i) the maximum number of alignments to display for each query-proteome pair; (ii) alignment metadata, such as query coverage, identity, and similarity; and (iii) target protein metadata, like target sequence size, annotation, and genomic co-localization of hits from different queries. The genomic co-localization filter is particularly valuable for identifying homologs of proteins whose genes are organized in operons. Filters can be applied globally to all results or defined on a per-query basis. For slider-based filters, the chart updates in real time, remaining responsive even when displaying over 10,000 hits. This reactivity enables users to fine-tune parameters to suit their research context, avoiding abrupt threshold effects. The control panel has a fixed position for real-time visualization of parameter changes, even when scrolling through the chart. It can also be collapsed to maximize the visualization area.

The Hit Graph provides a highly interactive and customizable overview of alignment results, plotting query sequences on the X-axis against proteomes on the Y-axis. By default, hits are represented as points that reflect their coverage percentage on the query sequence, with results grouped by genome on a single line. This design allows users to visually screen more than 50 proteomes on a single screen without scrolling. Hits for a protein within a specific proteome create distinct visual signatures, enabling straightforward comparison. With genomes ordered by taxonomic proximity, taxonomic prevalence patterns become immediately recognizable. In addition, a blast-like legacy display option shows hits based on their position on the query sequence for detailed alignment visualization. Interactive graphical features enhance usability, including Taxonomic Navigation (to navigate through taxonomic groups effortlessly), Graphical Hit Selection (to select specific hits directly on the graph), Genome Zoom (to focus on individual genomes for detailed inspection), and Hover Details (to view detailed information about specific hits by hovering over them). In short, this chart combines visual clarity with powerful interactivity, streamlining the analysis of complex alignment data.

The Selection Table, located below the hit chart, displays metadata for selected alignments and their corresponding hit proteins. Users can highlight or remove hits directly from the table. This feature is particularly useful when filtering alone cannot fully refine results. By combining graphical hit selection with the interactive table, users can precisely curate peers of interest. Below the table, an export feature allows users to save their selections, search parameters, and filtering criteria, in various formats, facilitating continued analysis with other tools without requiring additional file manipulation.

The Alignment Detail Display Area, located at the bottom of the page, displays the raw output of the HMM search program for the highlighted hit, providing in-depth alignment information for detailed examination.

## 3. Implementation

WHOOPER is a web application built in Python for its server side. The web frontend uses the D3.js data visualization library (https://d3js.org) for handling data visualization and exploration features. WHOOPER has been implemented to perform processing either on a local system or on a computing cluster, managed by the SGE scheduler.

To facilitate the simple deployment of specific instances, tailored to unique study contexts, WHOOPER includes procedures and scripts to configure an instance from a genome list retrieved from the NCBI dataset portal.

## 4. Acknowledgements

We are grateful to the INRAE MIGALE bioinformatics facility (MIGALE, INRAE, 2020. Migale bioinformatics Facility, doi: 10.15454/1.5572390655343293E12) and genotoul bioinformatics platform Toulouse Occitanie (Bioinfo Genotoul, doi: 10.15454/1.5572369328961167E12) for providing help and computing and storage resources.

