## Supplementary Figure 1 for "WHOOPER Web application for Hands-On identification of proteins co-Occurrence among Phyla, focused on user ERgonomics"

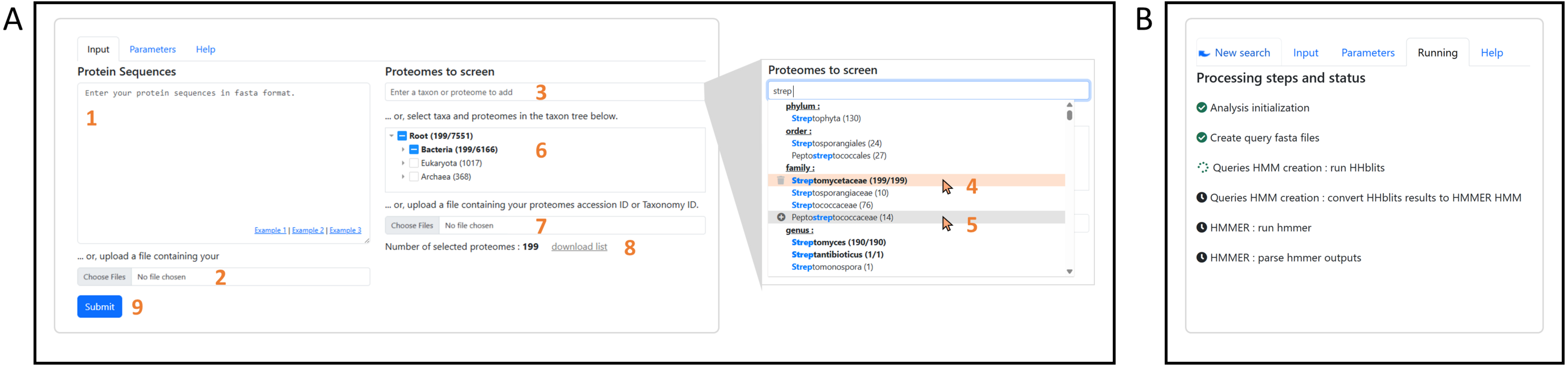


**Supplementary figure 1:** WHOOPER interfaces for launching and monitoring a search. (A) Submission interface. (B) Tracking interface. The search configuration process consists of importing query sequences for alignment and selecting the relevant proteomes for screening. Query sequences in FASTA format can be provided either through copy-pasting (1) or by uploading a file (2). Since selecting the appropriate proteomes is a key step in the analysis, various functionalities have been implemented to facilitate the selection of hundreds of proteomes by species or entire taxonomic groups: an autocomplete list (3) for adding (4) or removing (5) proteomes or taxa from the selection, an interactive taxonomic tree (6), and import/export options for accession and taxon ID lists(7,8). The latter feature facilitates easy iteration of new searches from previous analysis results. Once the search form is submitted (9), users are redirected to the tracking interface(B), which provides real-time updates on the analysis status and each step: queued, processing, completed, error. Any errors are displayed at the relevant step.
